# Improved short-channel regression for mapping resting-state functional connectivity networks using functional near-infrared spectroscopy

**DOI:** 10.1101/2023.06.12.543244

**Authors:** S. L. Novi, A. Abdalmalak, K. Kazazian, L. Norton, D. B. Debicki, R. C. Mesquita, A. M. Owen

**Affiliations:** Department of Physiology and Pharmacology, Western University, London, ON, Canada; Western Institute for Neuroscience, Western University, London, ON, Canada; Department of Psychology, King’s University College at Western University, London, ON, Canada; Clinical Neurological Sciences, Western University, London, ON, Canada; “Gleb Wataghin” Institute of Physics, University of Campinas, Campinas, SP, Brazil; Department of Psychology, Western University, London, ON, Canada

## Abstract

Resting-state functional connectivity (rsFC) is an attractive biomarker of brain function that can vary with brain injury. The simplicity of resting-state protocols coupled with the main features of functional near-infrared spectroscopy (fNIRS), such as portability and versatility, can facilitate the monitoring of unresponsive patients in acute settings at the bedside. However, accurately mapping rsFC networks is challenging due to signal contamination from non-neural components, such as scalp hemodynamics and systemic physiology. Physiological noise may be mitigated through the use of short channels which may be able to provide sufficient information to eliminate the need for additional measurement devices, decreasing the complexity of the experimental setup. To this end, we examined the extent to which systemic physiology is embedded in the short-channel data and improved short-channel regression to account for temporal heterogeneity in the scalp hemodynamics. Our findings indicate that using temporal shifts in the short-channel data increases the agreement, by 70% on average, between short-channel regression and regression that includes short channels and physiological recordings. Overall, this method decreases the need for additional physiological recordings when mapping rsFC networks, providing a viable alternative when such measurements are not available or feasible.

## 1. Introduction

Resting-state functional connectivity (rsFC) studies measure spontaneous brain activity while participants are instructed to remain still, close their eyes, and try to not focus on a particular thought (Biswal et al., 1997; Mesquita et al., 2010; Novi et al., 2016; Sasai et al., 2012, 2011). Functional magnetic resonance imaging (fMRI) studies have reliably documented a variety of rsFC maps in healthy controls (Beckmann et al., 2005; Damoiseaux et al., 2006; de Luca et al., 2006, 2005; van den Heuvel and Hulshoff Pol, 2010). These rsFC maps represent spontaneous low-frequency oscillations (SLFO) of the BOLD signal (usually < 0.15 Hz) that synchronize across specific brain regions and are thought to underlie distinct neural networks (Greicius et al., 2009; Sasai et al., 2012, 2011; Tong et al., 2019). Moreover, disruptions of rsFC have been shown to be clinically useful for diagnosing and monitoring patients with brain disorders, such as Alzheimer’s disease (Vemuri et al., 2012) and disorders of consciousness (Abdalmalak et al., 2021).

Functional near-infrared spectroscopy (fNIRS) is an alternative neuroimaging technique that also provides a proxy for neural activity through neurovascular coupling (Huppert et al., 2009b, 2007, 2006; Mesquita et al., 2009). FNIRS may be a superior alternative for fMRI as it is portable and does not pose any significant safety risks, which makes it ideal for monitoring patients at the bedside (Abdalmalak et al., 2017; Ayaz et al., 2022; Forero et al., 2017; Novi et al., 2023). The simplicity of resting-state measurements combined with the versatility of fNIRS makes it possible to monitor unresponsive patients, opening new avenues for assessing brain injured patients within the first days of hospitalization (Ayaz et al., 2022; Bicciato et al., 2022; Kazazian et al., 2021). However, extracting robust fNIRS-based rsFC maps is challenging due to contamination of the signal from non-neural components, such as scalp hemodynamics and systemic physiology (Abdalmalak et al., 2022; Bonilauri et al., 2021; Caldwell et al., 2016; Gagnon et al., 2011; Mesquita et al., 2010; Nasseri et al., 2018; Pinti et al., 2019; Scholkmann et al., 2022, 2013; Wyser et al., 2020; Yücel et al., 2021).

The non-neural components of the fNIRS signal are intrinsically related to the physical principles of the technique. More specifically, by shining and detecting near-infrared (NIR) light on the surface of the scalp, fNIRS estimates oxy-hemoglobin (HbO) and deoxy-hemoglobin (HbR) concentration changes as a proxy for local neural activity in the cortex (Boas et al., 2014; Scholkmann et al., 2014). However, in addition to the cortex, the NIR light interacts with superficial layers that are highly vascularized; hemodynamic changes in these layers yield spurious responses that confound the HbO and HbR changes that are due to neural activity (Saager and Berger, 2008, 2005). Roughly 94% of the signal from a regular fNIRS channel (i.e., source-detector (SD) distance around 3cm) comes from non-neural tissue, mainly the scalp vasculature (Brigadoi and Cooper, 2015). In addition, fluctuations in heart rate (HR), blood pressure, and breathing also induce hemodynamic changes in the brain vasculature that are independent of brain activity. That is to say, although these systemic hemodynamic fluctuations are physically present in the cortex, they do not reflect spontaneous cortical activity (Abdalmalak et al., 2022; McKetton et al., 2021; Scholkmann et al., 2022; Yücel et al., 2016). In resting-state studies, extra-cortical and systemic physiological noise contaminate rsFC networks and lead to high intra- and inter-participant variability (Abdalmalak et al., 2022). Removal of these non-neural contaminants is imperative for mapping reliable and meaningful fNIRS-based rsFC networks.

Additional measurements with short channels (SCs) and systemic physiology are currently the optimal methodological approach for removing non-neural components from the fNIRS signal (Abdalmalak et al., 2022; Caldwell et al., 2016; Scholkmann et al., 2022). SCs are channels in which the distance between sources and detectors are approximately 1 cm. Due to the diffusive character of NIR-light propagation in biological tissue, SCs are only sensitive to hemodynamic changes in the superficial layers of the head, mainly the scalp (Brigadoi and Cooper, 2015; Saager and Berger, 2008, 2005). Therefore, both short channels and systemic physiological measurements can be used to regress out non-neural components from the fNIRS signal. This decontamination is often performed within the General Linear Model (GLM) framework, in which these additional measurements are used as nuisance regressors of non-interest (Abdalmalak et al., 2022; Pinti et al., 2019; Santosa et al., 2020; Wyser et al., 2020). Recently, we have demonstrated that by regressing SCs, mean arterial pressure (MAP), HR, and CO2, fNIRS can reliably reproduce many of the rsFC networks that have been identified using fMRI, including the sensorimotor, auditory, and frontoparietal control networks (Abdalmalak et al., 2022). However, measurements of systemic physiology require external devices, which add complexity to the experimental setup and therefore limit fNIRS versatility and applicability.

As systemic physiology is not constrained locally and affects the hemodynamics of the head in the cortical and extra-cortical layers (Tong et al., 2019, 2012; Tong and Frederick, 2010; Yücel et al., 2015), we hypothesized that SCs should contain enough physiological information to avoid the need for additional physiological measurements completely, thereby decreasing the complexity of fNIRS experimental setups. To test our hypothesis, we investigated the extent to which systemic physiology is embedded in the short channels and then explored whether an improved short-channel regression could maximize the removal of non-neural components from the regular fNIRS channels.

## 2. Materials and Methods

### 2.1 Subjects and Experimental Protocol

The Research Ethics Board at Western University approved the study, which complies with the guidelines of the Tri-Council Policy Statement (TCPS): Ethical Conduct for Research Involving Humans. Before enrolling in the study, each participant provided written informed consent.

We acquired fNIRS resting-state data from 25 healthy young participants (13 females) with no history of brain disorders and with ages ranging from 20 to 31 years [mean (standard deviation) = 25 (4)]. Participants were seated in a comfortable chair and were instructed to remain still, close their eyes, and to not focus their thoughts on anything specifically. The duration of data acquisition lasted between 7 and 13 minutes. For a sub-cohort of 10 participants, additional measurements of HR, MAP, and CO2 (end-tidal capnograph waveform) were performed continuously and at the same time as the fNIRS acquisition (see Section 2.3). After each session, each participant was asked if they fell asleep. None of them reported falling asleep at any time during the experiment.

### 2.2 fNIRS Signal Acquisition

The fNIRS data were acquired with a commercial continuous-wave system (NIRScout, NIRx Medical Systems) at 3.9 Hz. The system works with 32 sources and 39 detectors. Each source is composed of 4 lasers centred at 785, 808, 830, and 850nm. Our optical probe allowed 121 long (SD distances around 3 cm) and 8 short channels (SD distance of 0.8 cm). The optical probe was hemispherically symmetrical and covered most of the head, including the frontal, temporal, parietal, and part of the occipital lobe. There were two short-channels in the frontal lobe, one in the temporal lobe, and one in the parietal lobe of each hemisphere. Figure 1 provides further details.

**Figure 1.**
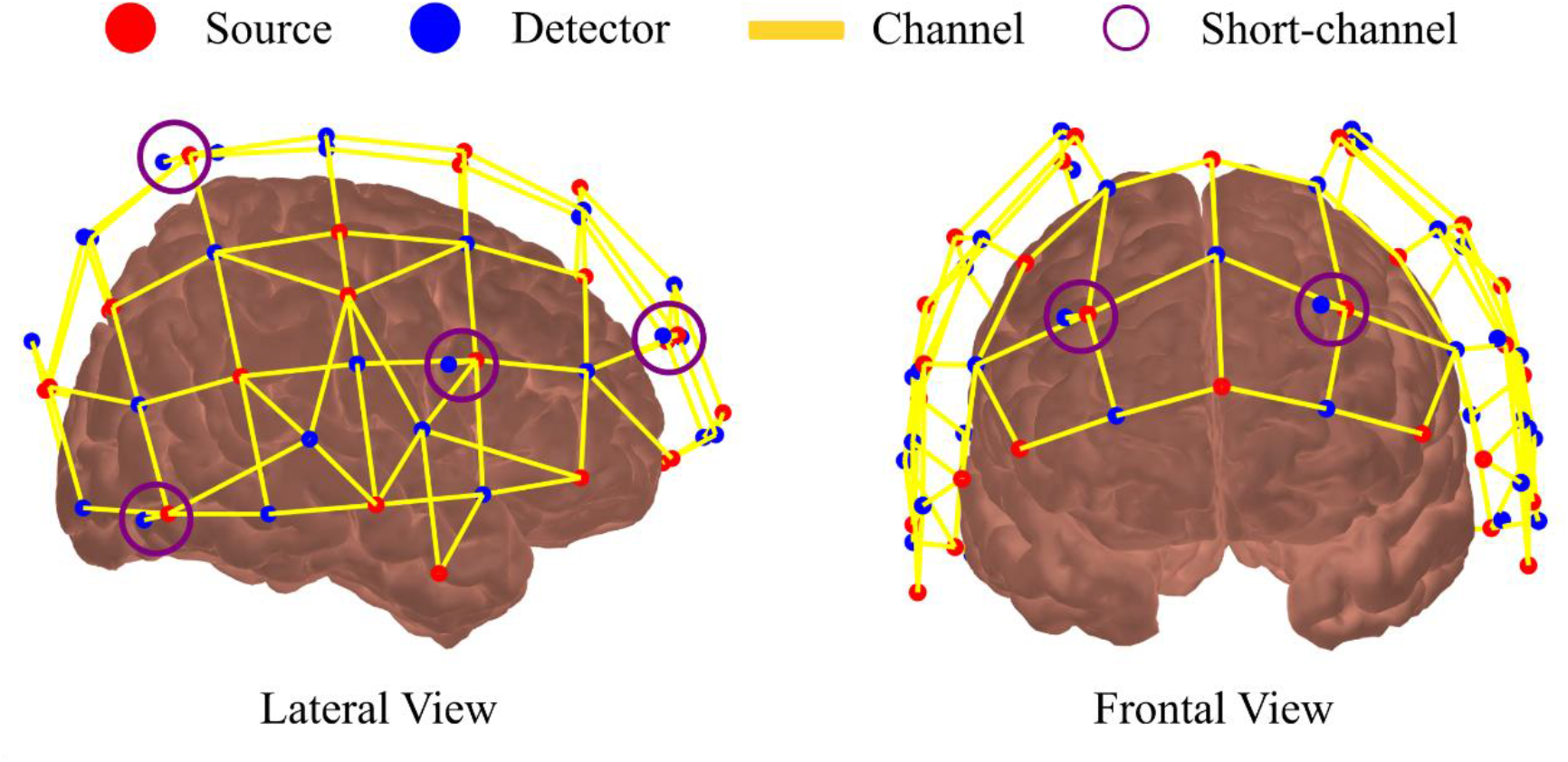
Lateral and frontal views of the optical probe used in this study. The optical probe was designed with 32 sources (red balls) and 39 detectors (blue balls). Our configuration allowed 121 regular channels (yellow lines) and 8 short channels (violet circles). Each short channel had a source-detector separation of 0.8 cm, and each regular channel had a source-detector separation of around 3 cm.

### 2.3 Physiological Recordings and Preprocessing

For a sub-cohort of 10 participants (Phys-dataset), additional systemic physiology was recorded at the same time as the fNIRS signal. HR and MAP were assessed with a Finometer Pro (Finapres Medical System, the Netherlands) with an acquisition frequency of 200 Hz, and CO_2_ was recorded with a capnograph (Oxigraph, Inc., United States) with an acquisition frequency of 4Hz. After calibration and synchronization of the two systems with the fNIRS device, 12 minutes of resting-state data was analyzed. The physiological measurements were resampled to the same frequency as the fNIRS data (3.9 Hz) then band-pass filtered between 0.009 and 0.08 Hz to be used as regressors of systemic physiology (see Section 2.5). The Phys-dataset has been previously published (Abdalmalak et al., 2022).

### 2.4 fNIRS Preprocessing

For preprocessing and analyzing the fNIRS data, we employed a MatLab toolbox developed in house and based on Homer2 functions (Huppert et al., 2009a), and followed the same procedure used by Abdalmalak et al., (2022). Briefly, regular channels with a signal-to-noise ratio (SNR) of less than 8 were discarded from further analysis (Forero et al., 2017; Novi et al., 2020a, 2020b). The SNR was defined as the average light intensity of a given channel divided by its standard deviation. To assess the quality of the SCs, we inspected their power spectrum individually and for each wavelength. SCs that lacked clear heart-rate peak (∼ 1Hz) or pink noise (1/f decay) in the frequency domain were discarded. One participant was removed for not having at least one good short channel.

Next, the light intensity of each remaining channel was converted to optical density, and motion artifacts were corrected with spline interpolation followed by wavelet decomposition (Novi et al., 2020b). Hemoglobin concentration changes were estimated for each channel using the Modified Beer-Lambert law with a differential pathlength factor equal to 6. The hemoglobin time series were band-pass filtered between 0.009 and 0.08 Hz for focusing on SLFO (White et al., 2009; Mesquita et al., 2010; Sasai et al.,2012; Niu et al., 2013). After band-pass filtering, non-neural components were removed from the fNIRS signals within the GLM framework by varying the design matrix accordingly.

### 2.5 Removal of Non-Neural Components under the GLM framework

Non-neural components were removed from the regular fNIRS channels within the GLM framework. In this framework, HbO and HbR of each regular channel (*Y*_*Hbx*_) is written as a linear combination of explanatory variables (design matrix **X)**, which are nuisance regressors in the case of resting-state data. Hence,

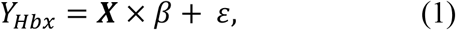

in which *β* is a vector, and *ε* is the error term. In the present work, the available explanatory variables are the short channels (**X**_SC_) and the additional measurements of HR, MAP, and CO_2_ (**X**_Phys_). We investigated how four GLM models performed when removing spurious correlations: (1) SC regression, which uses only the short-channel signals in the design matrix (**X** = **X**_SC_), (2) time-varying SC regression, which uses only the short-channel data but allows a temporal shift in the regressors (see section 2.5.1 for details), (3) physiology regression, which uses only the additional systemic physiological signals in the design matrix (**X** = **X**_Phys_), and (4) SC and physiology regression (SC+Phys), which combines the short-channel data without any temporal shift and additional systemic physiological signals in the design matrix (**X** = [**X**_SC_,**X**_Phys_]). The **X**_SC_ was built with the HbO and HbR time series of all good short-channels, and the **X**_Phys_ used MAP, HR, and CO_2_ (Abdalmalak et al., 2022). When regressing of MAP, HR and CO_2_, a maximum time-shift of 20 seconds (advancing or delaying) for each physiological regressor was allowed to account for the transit time between the physiological measurements (acquired from the finger and nose) and the fNIRS signal acquired in the head. The shift was defined as the one that maximized the correlation between the regular fNIRS channel and the regressor (Abdalmalak et al., 2022).

#### 2.5.1 Time-Varying Short-channel Regression (SC+Shift Regression)

Previous studies have shown that scalp hemodynamics follow spatially heterogenous patterns, which means that their hemodynamics are slightly different across the head (Gagnon et al., 2014, 2012; Goodwin et al., 2014; Kirilina et al., 2013, 2012; Wyser et al., 2020; Zhang et al., 2015). For resting-state data, most of this heterogeneity arises from delayed systemic physiology across the scalp due to its transit time (Wyser et al., 2020). This phenomenon has also been observed in the cortex in fMRI studies (Tong et al., 2019, 2012; Tong and Frederick, 2010).

In the present study, we investigated whether a simple temporal shift in the SC data that accounts for the heterogenous scalp hemodynamics could compensate for the absence of additional systemic physiological recordings. The method involves first performing a Principal Component Analysis n the **X**_SC_ matrix, then allowing a temporal shift (maximum ±10s) in each Principal Component (PC) that maximizes the correlation of each PC with the regular fNIRS channel (Zhang et al., 2015). In other words, **X**_SC_ is replaced by 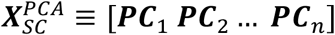 in which ***PC***_*i*_ represents the PCs of **X**_SC_, *n* is equal to the number of original regressors (*i.e*., all PCs are used), and a temporal shift is allowed for each PC. The PCs are shifted instead of the original channels to maximize the available temporal information as the PCs are not correlated across each other while the SCs are.

### 2.6 Correlation analysis and seed-based rsFC networks

After removing non-neural components, the temporal autocorrelation from HbO and HbR were removed with an autoregressive model, and HbT was computed as the sum of HbO and HbR (Arbabshirani et al., 2014; Barker et al., 2013; Santosa et al., 2017). The order of the autoregressive model was estimated for each channel as well as HbO and HbR with the Bayesian Information Criteria (Barker et al., 2013, 2016). This method has been shown to not disrupt the anticorrelation between HbO and HbR (Novi et al., 2023). After the removal of autocorrelation, functional connectivity matrices were computed independently for HbO, HbR, and HbT by computing the Pearson Correlation coefficient across the fNIRS channels.

To extract seed-based rsFC networks for each method, the individual correlation matrices were transformed to z-score coefficients by employing Fisher’s transformation. The Z-scores were averaged across all participants, yielding an average connectivity matrix for HbO, HbR, and HbT. Next, the sensorimotor, auditory, and fronto-parietal control (FPC) networks were identified by selecting the line of the average connectivity matrix that corresponded to a pre-defined seed (channel) for each network. The seeds (with respective networks) were located at: left primary motor cortex (sensorimotor network), left superior temporal gyrus (auditory network), and left middle frontal gyrus (FPC network).

### 2.7 Performance evaluation

In addition to extracting rsFC networks, the effect of allowing temporal shifts in the short-channel data was assessed by evaluating the correlation distributions (upper diagonal of the correlation matrix) at the group and individual levels for the Phys-dataset. We contrasted the standard and time-varying SC regressions against the SC+Phys regression by treating the latter as the gold standard. At the individual level, the comparison was quantified by the average sum of the absolute difference of each distribution:

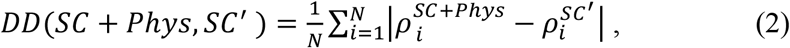

where *ρ*^*SC+Phys*^ is the correlation distribution for the SC+Phys regression, and 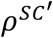 is the SC or SC+Shift regression. The index *i* represents each bin of the correlation distribution. Here, we computed *DD* for the entire distribution and for positive values only; a lower *DD* indicates a higher similarity between distributions. The *DD* values across methodologies were statistically tested with a two-sided Wilcoxon rank-sum test. At the group level, individual correlation distributions were concatenated and characterized by their mean and variance.

### 2.8 Estimation of mean arterial pressure and CO_2_ with short-channel data

To investigate how much systemic physiology is embedded in the short channels, we assessed the extent to which SC data could predict MAP and CO_2_ fluctuations. To do so, measured MAP and CO_2_ were modeled as a linear combination of each short channel:

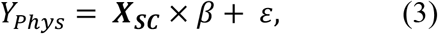

in which *Y*_*Phys*_ was the acquired preprocessed MAP and CO_2_ measurements. ***X***_***SC***_ was built with the HbO and HbR time-series of all good short-channels. A shift that maximized the correlation between each regressor and the physiological parameter was allowed. For this exploratory analysis, we varied the maximum allowed shift from 0 to 25 seconds. We used the bandpass-filtered data to focus on the frequency of interest of rsFC studies (i.e., between 0.009 and 0.08 Hz).

### 2.9 Estimation of heart rate with fNIRS data

As a complementary analysis, we estimated HR from fNIRS data using a regular channel because it usually has a higher SNR than short channels.

For extracting HR fluctuation, we employed a sliding window of 10 seconds that moved one frame per step to segment the HbO time series (prior to the bandpass filter) of a single channel. Next, we performed a Fourier transformation on each segment and defined the HR frequency as the maximum peak between 0.8 and 1.5 Hz, creating an estimated HR time series. Finally, the estimated HR was bandpass filtered between 0.009 and 0.08Hz and compared to the acquired HR with the Pearson Correlation (after removal of the autocorrelation).

## 3. Results

### 3.1 Physiology embedded in the fNIRS data

Before attempting to improve SC regression, we investigated how much systemic physiology is embedded in the fNIRS measurements. We found that fNIRS (regular channel) data could be used to accurately estimate externally measured HR data. Specifically, the correlations between the estimated and acquired HR for each participant ranged between 0.58 and 0.88, average of 0.76.

MAP and CO_2_ were modeled as a linear combination of short channels with the underlying assumption that both physiologies could be predicted if their effects were highly pronounced in the short channels. It was found that MAP could be estimated utilizing just the short channels when the maximum shift exceeded 4 seconds (Figure 2A). Above this threshold, the quality of estimated MAP did not depend on the maximum allowed shift. In fact, MAP from 8 out of 10 participants could be estimated with correlations above 0.5, which is a good agreement given that the autocorrelation was removed from both time series. The two participants with correlations below 0.5 had only 1 and 2 good short channels, respectively, and two of the three participants with correlations higher or equal to 0.7 had 5 good SCs, suggesting that the number of good short channels plays a crucial role in MAP estimation. Overall, our results indicate that a great part of the MAP signal in the low-frequency range is embedded in the short channels and can be estimated from them if enough good short channels are available.

**Figure 2.**
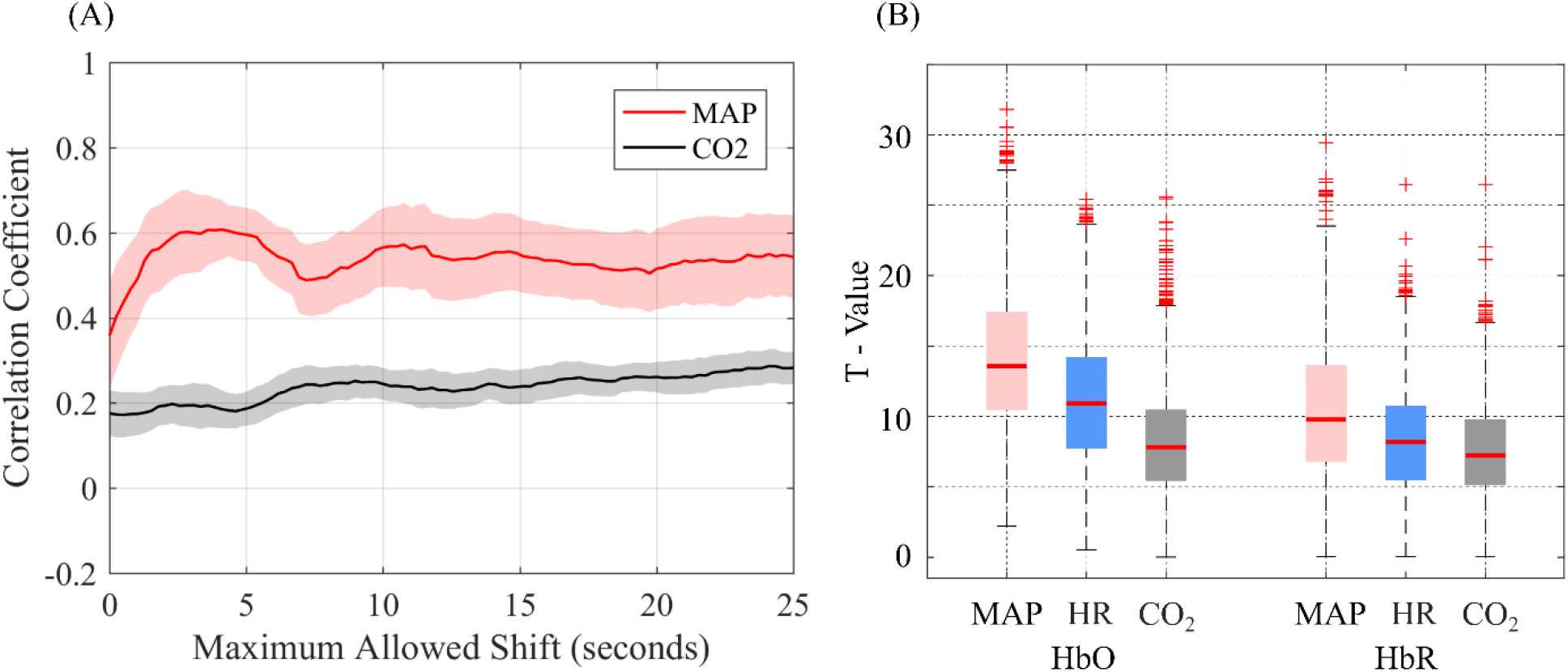
(A) Average Pearson correlation coefficient between estimated and measured MAP and CO_2_ for the maximum allowed shifts varying from 0 to 25 seconds. The shaded region is the standard deviation across participants. There was a total of 10 participants. (B) Each boxplot depicts the distribution of absolute T-values obtained from the betas estimated for each regressor in the Phys Regression for HbO and HbR: MAP in red, HR in blue, and CO_2_ in gray. Each distribution contains 1177 points. The T-value is proportional to the importance of each regressor.

In contrast, we found that the CO_2_ measurements could not be predicted by simply shifting the information in the SCs (gray curve in Figure 2A). At the low-frequency band investigated, the prediction was poor for any time shift, indicating that this is not due to asynchrony from slow transit times. These findings suggests that low-frequency hemodynamics of the scalp are not linearly affected by CO_2_ fluctuations during resting state.

As a complementary analysis, we attempted to estimate the effect of each physiological parameter when decontaminating the regular fNIRS channels. To this end, we extracted the T-values associated with MAP, HR, and CO_2_ when physiology was regressed from the long channels (*i.e*., equation (1) with **X** = [**X**_Phys_]). Figure 2B shows the absolute T-value for each regressor for all regular channels in all participants. Although there is high variability in the absolute T-values associated with each physiological parameter, MAP and CO2 present the highest and lowest T-values, respectively. As the T-distribution has an exponential decay towards higher absolute values, additional measurements of MAP can explain the fluctuations in the typical fNIRS channels much better than CO2 recordings in the periphery. Thus, the physiological information embedded in the SC data, which is mainly MAP, might be sufficient to properly decontaminate the fNIRS signal and extract meaningful rsFC networks.

### 3.2 SC+Shift removes more spurious correlations than the standard SC regression at the group level

To explore whether SC regression could be improved without the need for additional physiological measurements we investigated whether shifts in the SC data would perform equivalently to SC+Phys regression. Overall, for HbO and HbT, shifts in the short channels led to a higher agreement with the SC+Phys regression than SC alone (Figure 3). The shifts decreased the standard deviation (inferred with a Gaussian fit) of the correlation distributions for HbO, HbR, and HbT(see Table 1) demonstrating that the SC+Shift approach removes more spurious correlations and has a better agreement with the SC+Phys regression than the standard SC regression.

**Figure 3.**
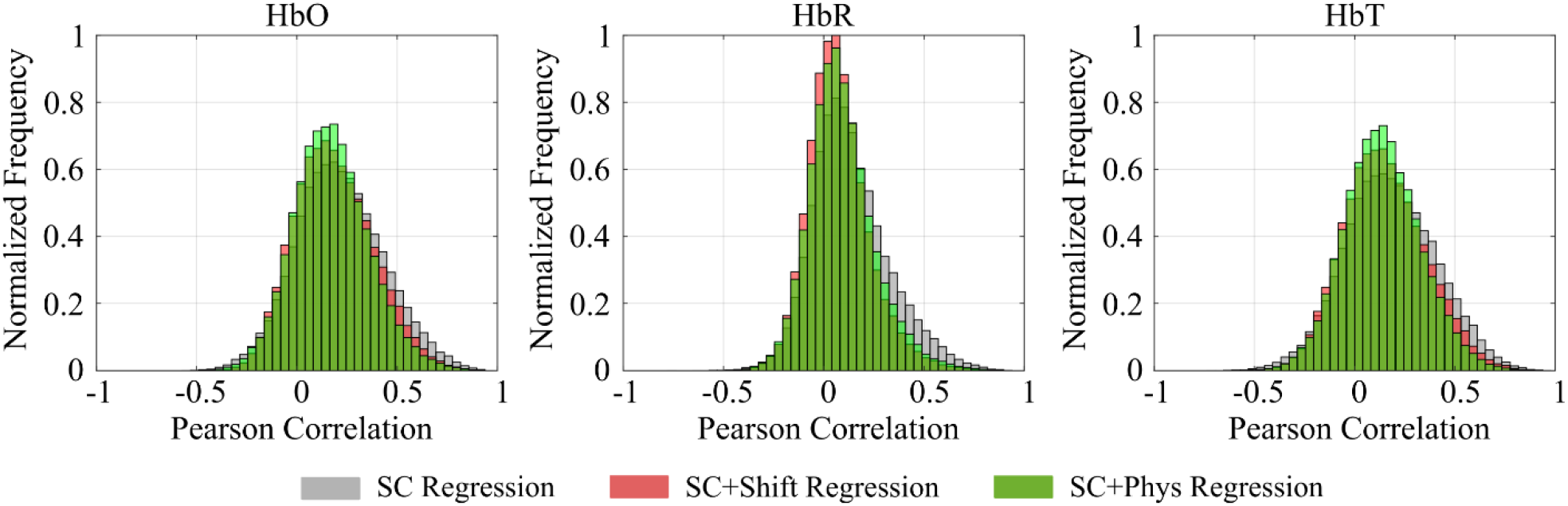
Group correlation distributions for HbO, HbR, and HbT, extracted with the Phys dataset. The gray, red, and green are the distributions extracted after the SC, SC+Shift, and SC+Phys regressions, respectively. The SC+Shift has a higher agreement with the SC+Phys methodology.

**Table 1.**
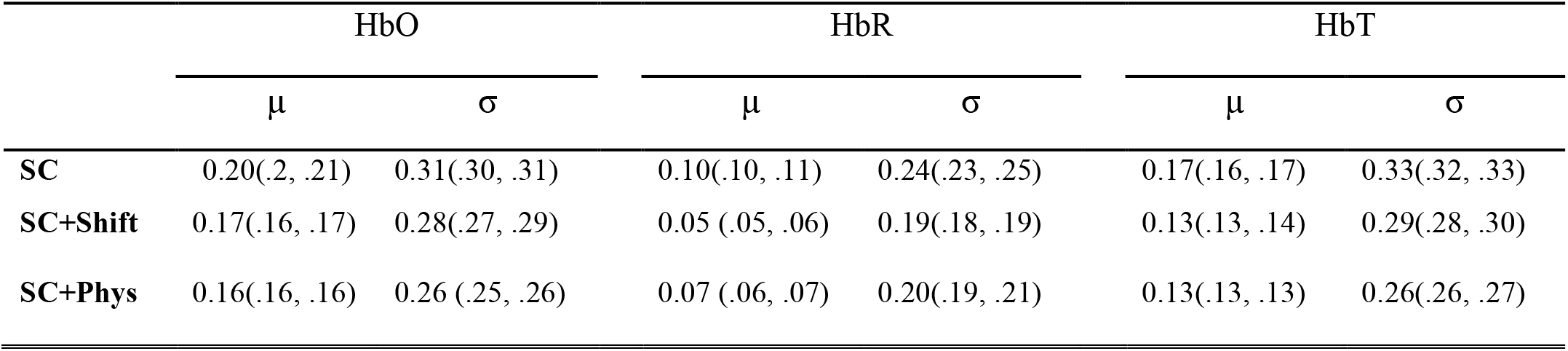
Estimates after modeling the data shown in Figure 3 with a Gaussian distribution with mean (µ) and standard deviation (σ). Values inside the parenthesis show the 95% confidence intervals of the estimated parameters. Overall, the SC+Shift has a higher agreement with the SC+Phys than the standard SC regression.

In addition to the correlation distributions, we investigated common rsFC networks extracted for the group with the seed-based approach for HbT. We focused on HbT because it has been shown to yield the highest reproducibility and agreement with fMRI when compared with HbO and HbR (Abdalmalak et al., 2022; Culver et al., 2005; Novi et al., 2023, 2016; Sheth, 2004). Figure 4 shows the sensorimotor, auditory, and FPC networks acquired with all preprocessing methodologies. As the removal of non-neural components becomes more effective, the rsFC becomes more localized and in better agreement with the fMRI literature. For example, the sensorimotor network covered all probed brain regions without regression, while it became limited to the sensorimotor cortex when both scalp and systemic physiology were removed (SC+Phys regression). Shifting the short-channel data localized the sensorimotor network and increased the agreement between it and the SC+Phys regression compared to the standard SC regression. In fact, the standard SC regression was not sufficient to remove all spurious correlations outside the sensorimotor cortex, such as in the temporal lobe, while the SC+Shift successfully did and maintained a strong correlation between the seed and its homotopic brain region.

**Figure 4.**
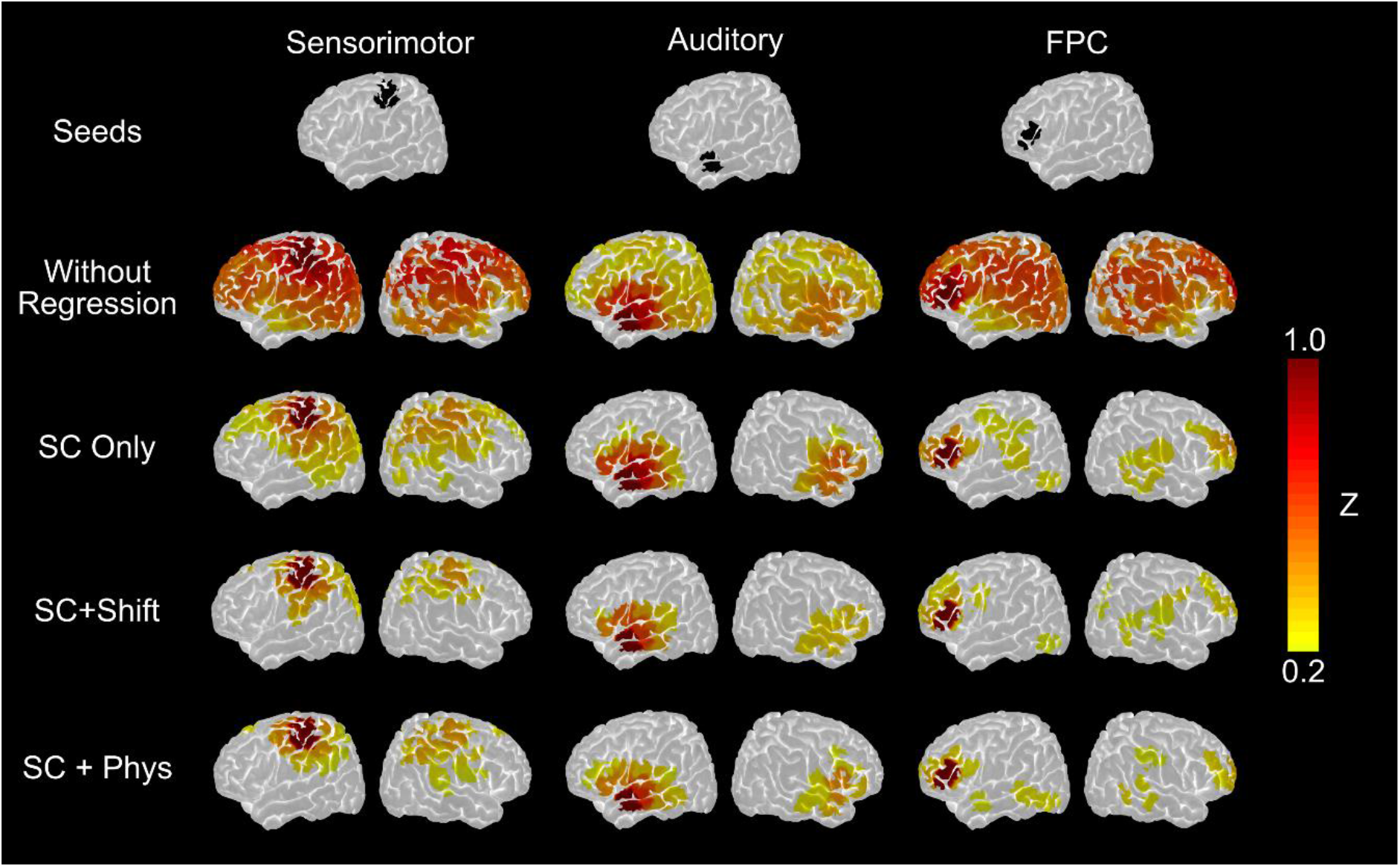
Sensorimotor, Auditory, and FPC seed-based networks extracted for HbT at the group level for the Phys dataset. Each column in the top row shows the location of each seed, which is an fNIRS regular channel. The following rows show the obtained networks after each method of regression, including the one without any regression. The networks become more localized with a higher agreement with the fMRI literature as the spurious correlations are regressed out. SC+Shift presents the highest agreement with SC+Phys regression.

Regarding the auditory and FPC networks, since the SC regression achieved results similar to the SC+Phys regression, we did not expect many improvements with the SC+Shift regression. To clarify, the auditory network was already very localized with the SC regression in such a way that the SC+Phys regression did not bring any additional advantages. However, it is important to note that despite the unnoticeable change achieved by shifting the short channels, the auditory network was not disrupted. For the FPC, the SC+Shift regression removed additional spurious correlations (compared to the SC regression) in regions that do not belong to the FPC network, mainly in the left primary motor cortex. Hence, for the group analysis, shifting the short-channel data is a viable and straightforward solution that improves the performance of standard SC regression when rsFC networks are not localized, and the shifts do not disrupt localized networks by removing important connections.

### 3.3 SC+Shift performed better than the SC regression at the individual level

After verifying the performance of SC+Shift at the group level, we inspected the results to see if they were consistent at the individual level. Figure 5 shows the HbT correlation distributions of each participant for each type of regression.

**Figure 5.**
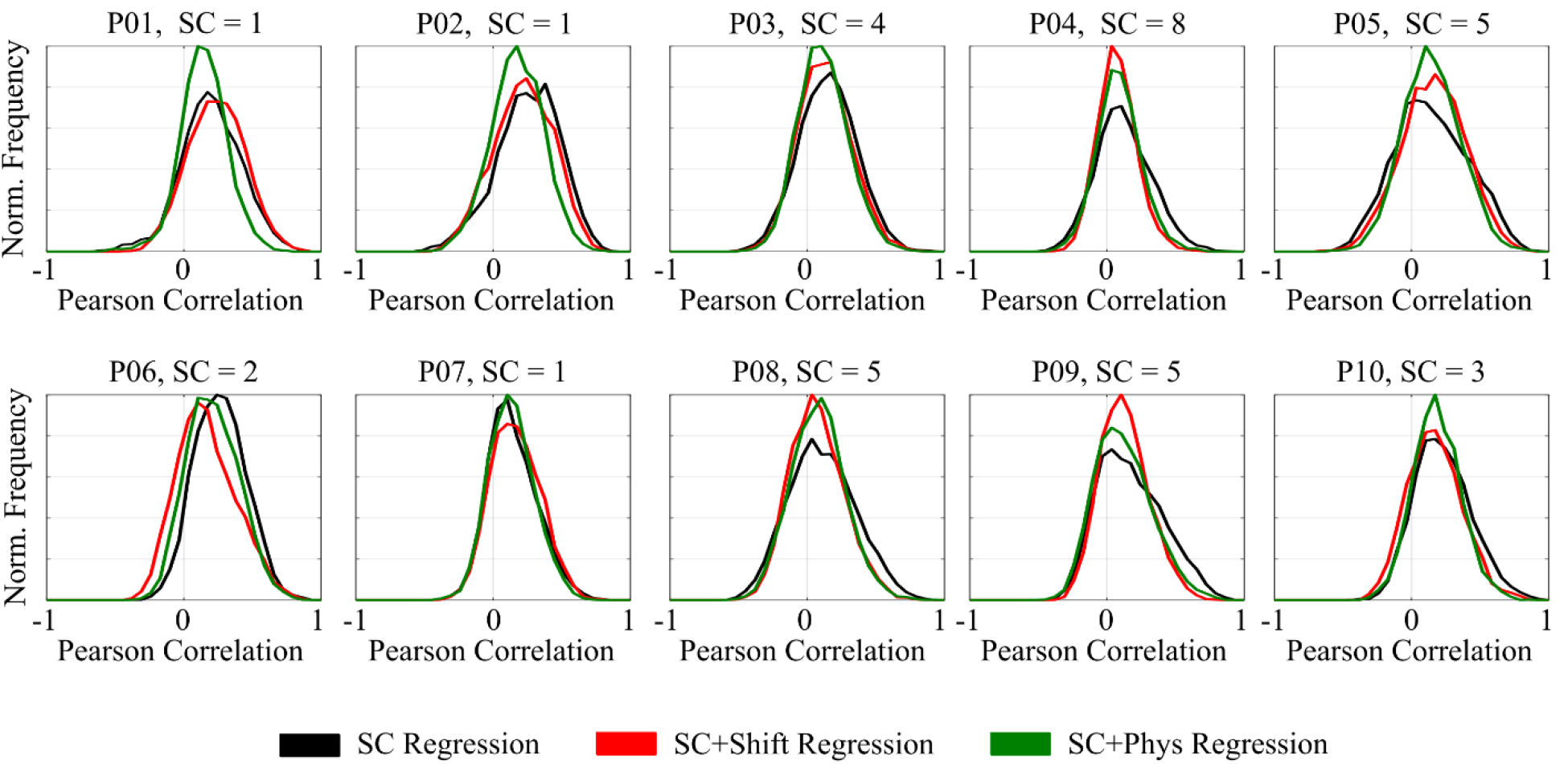
Individual correlation distributions for HbT extracted from the Phys-dataset after SC (black), SC+Shift (red), and SC+Phys (green) regressions. The title in each subplot shows the participant index and the number of good short channels available for that participant. At the individual level, the shift in the short-channel data presented a visible improvement in half of the participants. The SC+Shift improvement tends to be higher when more short channels are available.

By treating the SC+Phys as the gold standard, visual inspection indicates that the SC+Shift was very effective for 5 participants: P04, P05, P08, P09, and P10. For these participants, the agreement between the short-channel regression and SC+Phys became considerably higher when the shifts were allowed. Interestingly, these five participants are the ones that were most similar when estimated and measured MAP were compared, and all of them had at least three good short channels (3 participants had 5 good SCs, 1 had 8, and 1 had 3). Notably, P03 had four good short channels and presented a small improvement even though SC regression was already very similar to the SC+Phys regression.

Moreover, our quantitative analysis also showed that the shifts in the short-channels improved the removal of spurious correlations by significantly increasing the agreement with the SC+Phys regression for the entire distribution (Figure 6A) and for positive values (Figure 6B), which is often the correlation range of interest. The median normalized absolute distance *DD* decreased from 0.51 to 0.35 and from 0.47 to 0.27 for the total and positive values, respectively, which means an increase of 70% on average in the agreement between regressing the fNIRS signals with only the SC data and with the combination SC and additional physiological measurements. Overall, allowing shifts in the SC data increased the performance of the SC regression for participants that exhibited a poor standard SC regression, and, more importantly, the shifts did not negatively effect the correlation distributions for those who already had good agreement between SC and SC+Phys regressions.

**Figure 6.**
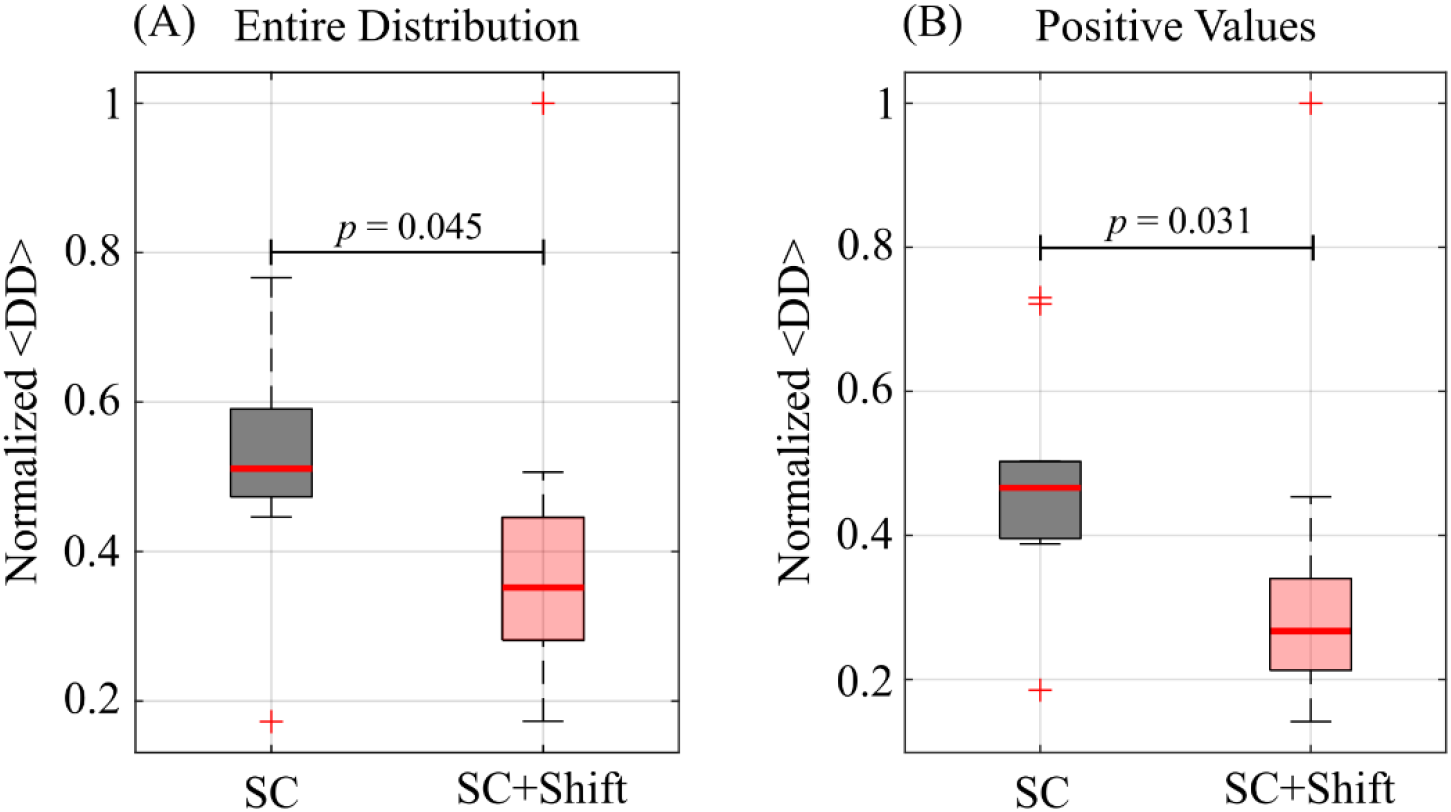
Normalized average sum of the absolute difference (<DD>, Equation 2) computed between the pairs of distributions (SC+Phys, SC) and (SC+Phys, SC+Shift) for the entire distribution (A) and only positive values (B). The normalization was computed within the condition by dividing *DD* by the highest value between *DD*(SC+Phys, SC) and *DD*(SC+Phys, SC+Shift). The number of points in each distribution is 10.

### 3.4 Improved network localization with SC+Shift on the full dataset

To further validate our findings, we applied the SC and SC+Shift regressions to the entire dataset of 24 participants. Figure 7 displays the group-level sensorimotor, auditory, and FPC networks extracted using HbT. Overall, the SC+Shift regression generated more localized sensorimotor networks than the standard SC regression. For example, the SC+Shift sensorimotor network resembles the SC+Phys network from the Phys-dataset (Figure 4), being constrained to the sensorimotor cortex. The SC+Shift FPC network was also better localized and showed stronger agreement with the SC+Phys FPC network displayed in Figure 4. The shift in the SC data helped to refine the networks while still preserving the homotopic connections, which are a hallmark of resting-state functional brain connectivity.

**Figure 7.**
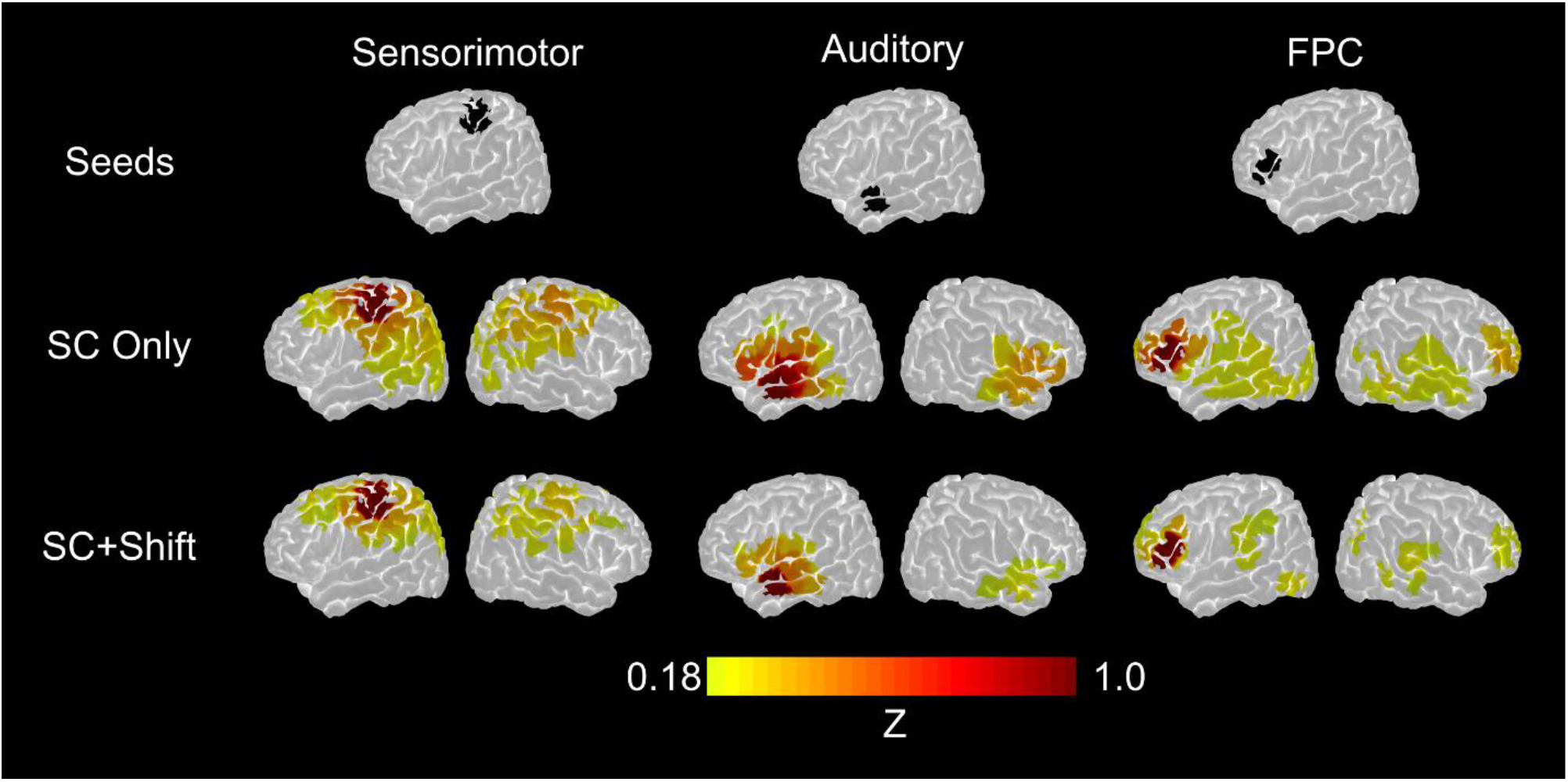
Sensorimotor, Auditory, and FPC seed-based networks extracted for HbT at the group level for all 24 participants. Each column in the top row shows the location of each seed, which is an fNIRS regular channel. The second and third rows show the rsFC networks obtained with the SC and SC+Shift methods, respectively. SC+Shift regression yields the most localized networks with good agreement with the fMRI literature.

## 4. Discussion

Resting-state functional connectivity is an attractive biomarker of brain function that can vary with brain injury. The simplicity of resting-state experiments combined with the versatility of fNIRS can facilitate the monitoring of patients in acute settings at the bedside. However, accurately mapping rsFC networks with fNIRS is challenging due to contamination from non-neural components, such as scalp hemodynamics and systemic physiology (Abdalmalak et al., 2022; Ayaz et al., 2022; Lanka et al., 2022; Mesquita et al., 2010; Scholkmann et al., 2022; Yücel et al., 2016). Recently, our group has shown that recording additional physiological parameters, such as MAP, HR, and CO_2_, can be used to effectively decontaminate the fNIRS signal (Abdalmalak et al., 2022). Here, we extend this previous work by demonstrating that short channels provide sufficient physiological information to decrease the need for additional measurements altogether. Through improvements in SC regression, using temporal shifts in the SC data to account for spatial heterogeneity in scalp hemodynamics, it is possible to decrease the need for additional physiological recordings when mapping rsFC networks, providing a viable alternative when physiology is not available.

When it is not possible to measure all physiological parameters, an often-asked question is which systemic physiology should be prioritized. This question has no unique answer because different experimental protocols will induce different physiological changes in different frequency bands (Scholkmann et al., 2022; Wyser et al., 2020). For example, while we found that in this resting state data CO_2_ contributed less to the SLFO signals than MAP and HR (Figure 2B), it has been shown to play a crucial role when speech-related stimuliare used (Scholkmann et al., 2013). While MAP is a well-known confounding factor in fNIRS data, HR effects are often overlooked on the basis that a simple low-pass filter can eliminate its contaminating influence (as HR frequency is around 1Hz for healthy adults). However, our results indicate that the low (0.04-0.15Hz) and very low-frequency (<0.04Hz) components of HR, associated with circadian rhythms, core body temperature, metabolism, hormones, and blood-pressure regulations (Shaffer et al., 2014), increase the covariance across fNIRS channels, resulting in inflated correlation distributions. Fortunately, SC regression can remove MAP and HR contributions, and its efficacy is maximized by allowing shifts in the SC data.

To investigate the extent to which information about systemic physiology is embedded in the hemodynamics of the scalp, MAP and CO_2_ were modelled as a linear combination of short channels. Overall, we found that MAP could be estimated with high precision, indicating that SCs can recover most of the information provided by external MAP recordings. However, the estimated MAP could not be used as a nuisance regressor in the GLM as its estimation required additional recordings with an external device. In other words, this approach would not decrease the experimental complexity involved in measuring additional physiological parameters. As an alternative, we opted to allow temporal shifts in the SC data instead of using the estimated parameters as regressors, which is similar to the way MAP was estimated and accounts for the spatial heterogeneity in the hemodynamics of the scalp.

A relevant methodological difference between estimating HR and the other parameters is that HR was extracted with a Fourier transformation without requiring external data. As the external HR measurements were only used to evaluate the quality of the estimations, the estimated HR could be used as a nuisance regressor. We tested this scenario and observed no clear improvements (results not shown) that would justify the addition of estimated HR in the SC+Shift method. This lack of improvement indicates that the estimated HR does not provide clear complementary information to the SC data when the shifts are allowed. In addition, we also explored the possibility of extracting a proxy to CO_2_ following the same approach we did for the HR, but focusing on the breathing frequency window (*i.e*., between 0.2 and 0.4 Hz instead of 0.8 and 1.5 Hz). With this approach, the correlations between the external and estimated CO_2_ were weak, confirming that CO_2_ effects are not as influential as HR and MAP effects for studying spontaneous low-frequency oscillations.

The main challenge in developing methodologies for fNIRS with real data is that the ground truth is often unknown. For task-based protocols, this problem can be overcome with semi-synthetic data in which a hemodynamic response function (HRF) is added to resting-state recordings. Each investigated method is then judged based on the number of false and true positives and by how similar the recovered HRF compares to the simulated one. For rsFC studies, although promising attempts have been made (Lanka et al., 2022), the community still lacks a standard or broadly accepted methodology to simulate data. Hence, additional caution must be taken to evaluate the performance of each investigated algorithm. For example, one cannot assume that the best method is the one that removes more correlations because there are algorithms mathematically built to remove covariance across channels independent of the origins of it, such as Principal Component Analysis. In this respect, one might criticize the present work with the argument that the SC+Shift regression is constructed in such a way that it should remove more correlation than the standard SC regression. However, it is important to note that we evaluated each method by verifying how each one compares to the SC+Phys regression. We previously demonstrated that the standard SC regression was not sufficient to reproduce some of the most common rsFC networks identified with fMRI. The addition of MAP, HR, and end-tidal CO2 not only improved the agreement between fNIRS and fMRI but also decreased the inter- and intra-subject variability (Abdalmalak et al., 2022). Therefore, the increase in similarity between SC+Shift with the SC+Phys regression (and consequently with the BOLD-fMRI) is a good quality measure for this specific case. Moreover, the SC+Shift regression has a solid physiological motivation that maximizes the removal of non-neural components by considering the origin and spatial heterogeneity of hemodynamics of the scalp.

The main assumption of using a GLM is that the signal of interest can be modeled as a linear combination of the explanatory variables. In the fNIRS context, this framework implies that only temporal patterns that are present in both the hemoglobin time-series from the regular channels and nuisance regressors can be removed. More specifically, the efficacy of the standard SC regression is dependent on the effect of systemic physiology on both cortical and extra-cortical vascular hemodynamics in terms of similarity and synchronization. We propose using an additional shift in the SC data after decomposing them into PCs. In doing so, we are effectively adding a transfer function to the GLM that accounts for the transit time between the effects of the systemic physiology in both the scalp and cortical tissue. The full meaning of the shift depends on several factors, such as the location of the short channels with respect to the regular channel. For example, for the scenario in which there is only one or two short channels, and the SCs are far from the regular channel, the shift will mostly account for transit time within the scalp tissue. The transit time has been reported to be up to 10 seconds on average (Zhang et al., 2015) for SLFO assessed by SCs in the scalp. In the case of several short channels, because more information is available, the shift may also account for the difference in transit time of systemic physiology between the scalp and the cortical tissue.

Finally, we would like to discuss that the method introduced here contributes to the broad and emerging field of systemic physiology augmented fNIRS (SPA-fNIRS; see Scholkmann et al., 2022). In general terms, SPA-fNIRS argues that systemic recordings are necessary to better interpret and understand fNIRS results. SPA-fNIRS can separate the neural from non-neural components of the signal by regressing out the measured physiology as we did with the SC+Phys regression. Although additional measurements minimize misleading results and conclusions, current SPA-fNIRS limits the fNIRS applications by adding complexity to the experimental setup, thereby decreasing fNIRS portability and versatility. With this in mind, we emphasize that alternative methods for decontaminating the fNIRS signal are imperative for fNIRS to reach its full potential and become a stand-alone technique. These advanced methods will play an important role, particularly, in protocols in which acquiring additional systemic physiology is not feasible.

## 5. Conclusion

In conclusion, the present work aimed to verify whether short-channel regression could be improved to avoid the need for acquiring additional systemic physiology. To this end, we investigated the role of MAP, HR, and CO_2_ in both regular and short channels. In the context of mapping rsFC networks, our results indicate that MAP is the most important physiological confounding factor and has a strong effect in both cortical and superficial tissue layers. Moreover, we found that allowing shifts in the short-channel data increases the agreement between SC regression and SC+Phys regression by 70% on average. We suggest that this methodological approach, which only requires short channels to remove non-neural components from the fNIRS signals, is a valuable alternative when additional recordings of physiology are not available or feasible.

## 8. Data Availability Statement

The data supporting the findings of this research are available on request to the corresponding author, pending a formal data sharing agreement and approval from the local ethics committee.

